# Cell–cell adhesion drives patterning in stratified epithelia

**DOI:** 10.1101/2023.11.24.567740

**Authors:** Yosuke Mai, Yasuaki Kobayashi, Hiroyuki Kitahata, Takashi Seo, Takuma Nohara, Sota Itamoto, Shoko Mai, Junichi Kumamoto, Masaharu Nagayama, Wataru Nishie, Hideyuki Ujiie, Ken Natsuga

## Abstract

Epithelia consist of proliferating and differentiating cells that often display patterned arrangements. However, the mechanism regulating these spatial arrangements remains unclear. Here, we show that cell–cell adhesion dictates multicellular patterning in stratified epithelia. When cultured keratinocytes, a type of epithelial cell in the skin, are subjected to starvation, they spontaneously develop a pattern characterized by areas of high and low cell density. Pharmacological and knockout experiments show that adherens junctions are essential for patterning, whereas mathematical modeling indicates that cell–cell adhesion alone is sufficient to form regions with high/low cell density. This phenomenon, called cell–cell adhesion-induced patterning (CAIP), influences cell differentiation and proliferation through Yes-associated protein modulation. Starvation, which induces CAIP, enhances the stratification of the epithelia. These findings highlight the intrinsic self-organizing property of epithelial cells and indicate that CAIP modulation might promote wound healing in clinical settings.

## Introduction

Epithelial tissue is composed of layers of cells that cover the surfaces of organs, and their homeostasis is maintained through the spatial organization of cells with various fates. Epithelial cells are distributed in a coordinated manner and typically exhibit visible patterns in stratified epithelia, such as human fingerprints^1^, bird feathers^2^, and mouse tail scales^3–5^, at the macroscopic level. Genetic, developmental, and environmental factors contribute to the unique arrangement of stem cells, proliferating cells, and differentiating cells in tissues, leading to context-dependent epithelial patterning.

Keratinocytes are the primary cells in the epidermis, the stratified epithelium of the skin. Previous studies have highlighted keratinocytes’ ability to develop patterns. Seminal research by Green and Thomas demonstrated that human keratinocytes formed patterns resembling human fingerprints when cultured on feeder cells^6^. Subsequent studies revealed that human keratinocytes can self-organize into clusters expressing stem cell markers^7,8^, implying that the multicellular arrangements might affect cell fate. However, the characteristics and mechanisms underlying keratinocyte patterning are not yet thoroughly understood.

Cell–cell adhesion is a characteristic of epithelia and is supported by specialized junctional complexes, including adherens junctions (AJs). AJs are composed of cadherins and catenins and provide mechanical attachments between neighboring epithelial cells^9^. Perinatal lethality has been observed in mice lacking any of the AJ components^10–13^, underscoring the significance of AJs.

Here, we show that cell–cell adhesion governs keratinocyte patterning. The patterning is mediated by AJs and facilitates cell dynamics with the help of the Yes-associated protein (YAP) pathway. Our findings elucidate the molecular and cellular basis underlying the spatial organization of cells in the epidermis.

## Results

### Spontaneous patterning of keratinocytes

First, we observed the morphology of epithelial cell sheets by employing HaCaT cells, an immortalized keratinocyte line^14^. The cells were seeded in a high-calcium medium containing 10% fetal bovine serum (FBS) and allowed to reach full confluence in one day (day 1; Fig. 1a). On day 4, the cells displayed a self-organized pattern, comprising areas of high and low cell density (Fig. 1a, b). HaCaT cells were originally derived from the back skin of a 62-year-old male and not from a single cell^14^; therefore, heterogeneous cells might have formed the regions of high or low cell densities. To rule out this possibility, we performed single-cell cloning of HaCaT cells by integrating a puromycin cassette and the Cas9 gene into the cells, the latter of which enabled further knockout experiments^15^. Single-cell-cloned HaCaT cells developed a self-organized pattern similar to that observed in the parental cells (Supplementary Fig. 1), indicating the presence of an intrinsic cell property that gives rise to regions of high or low cell density. Morphologically, phalloidin staining revealed that cells in the areas of high cell density were cuboidal, compact, and stratified, whereas those in the areas of low cell density were flat (Fig. 1c).

**Figure 1.**
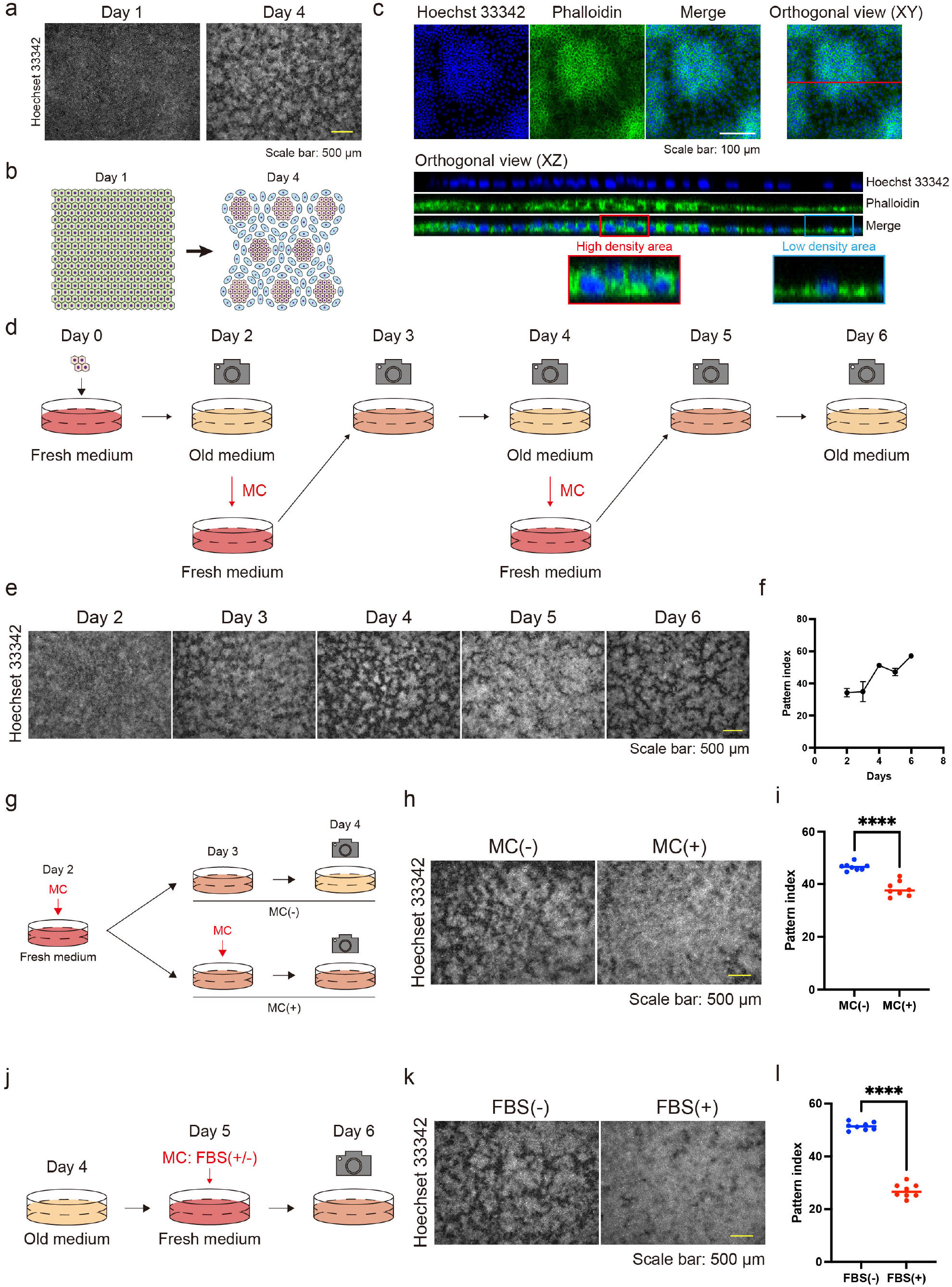
Serum starvation induces keratinocyte pattern formation. **a** Immunofluorescent images of keratinocytes displaying a self-organized pattern 1 day and 4 days after seeding. Nuclei are labeled with Hoechst 33342. Scale bar: 500 µm. **b** Schematic diagram of keratinocyte pattern, comprising areas of high and low cell density. **c** Orthogonal immunofluorescent images of keratinocyte pattern. Blue indicates nuclear labeling with Hoechst 33342. Green indicates actin labeling with phalloidin. The XZ orthogonal images correspond to the red line of the XY orthogonal view. The red and blue rectangles are enlarged to show a high-density area and a low-density area, respectively. Scale bar: 100 µm. **d** Schematic diagram of medium changes (MCs) and observation points. **e** Immunofluorescent images of keratinocytes following MCs. Nuclei are labeled with Hoechst 33342. Scale bar: 500 µm. **f** Pattern index during the cell culture. Data are presented as mean values ± standard error of the mean (SEM). N = 16 for each time point. **g** Schematic diagram of the experiments comparing keratinocyte pattern formation with or without MC. **h** Immunofluorescent images of keratinocytes with or without MC. Nuclei are labeled with Hoechst 33342. Scale bar: 500 µm. **i** Pattern index with or without MC. N = 8 for each group. **j** Schematic diagram of the experiments comparing keratinocyte patterns after MC with or without fetal bovine serum (FBS). **k** Immunofluorescent images of keratinocytes after MC with or without FBS. Nuclei are labeled with Hoechst 33342. Scale bar: 500 µm. **l** Pattern index after MC with or without FBS. For **i** and **l**, data are presented as mean values. Statistical analysis was performed with two-tailed Mann–Whitney U tests. ****, *P* < 0.0001.

Time-lapse images of keratinocytes during patterning revealed that cells initially moved randomly, and subsequently, the areas of high and low cell density developed spontaneously (Supplementary Fig. 2a, b; Supplementary Movie 1). This self-assembling pattern was obscured one day after a medium change (MC) (day 4 to day 5; Fig. 1d, e) and reappeared on day 6 (Fig. 1d, e). Time-lapse images showed that cells in the areas of high cell density migrated toward regions of low cell density after MCs and that the areas of high/low cell density reformed (Supplementary Fig. 3a– d; Supplementary Movie 2, 3). The pattern was quantified with an ImageJ plugin, in which the pattern index increases as the distinction between areas of high and low cell density becomes more evident (see Methods). This quantitative analysis confirmed that the pattern was disrupted by MCs (Fig. 1f).

### Serum starvation induces the self-organizing pattern of keratinocytes

The time course studies (Fig. 1d–f; Supplementary Movie 1–2; Supplementary Fig. 2–3) led us to speculate that the starvation of the culture medium induced the formation of keratinocyte patterns. To test this hypothesis, we compared groups with and without MCs and found that the pattern was disturbed in the group with MCs (Fig. 1g–i). The automated perfusion culture system (Supplementary Fig. 4a) demonstrated that the procedure of the MC itself did not significantly impact the phenotype, as the cells maintained at a low perfusion rate exhibited more pronounced patterns than those maintained at a high perfusion rate (Supplementary Fig. 4b, c). To further investigate the factors contributing to pattern formation in the culture medium, we compared cells that underwent MCs with or without 10% FBS (Fig. 1j–l). One day after the MC, the pattern disappeared in the FBS group but not in cells without FBS (Fig. 1j–l; Supplementary Fig. 5a, b; Supplementary Movie 4). These results indicate that serum starvation is crucial for keratinocyte patterning.

### Cell–cell adhesion through adherens junctions is essential for pattern development

To elucidate the underlying mechanism(s) behind the pattern formation, we performed RNA sequencing (RNA-seq) analysis to compare HaCaT cells cultured under high- or low-density conditions, as the pattern consisted of areas of high and low cell density. The gene ontology biological process terms enriched among the differentially expressed genes (DEGs) through DESeq2^16^ included cell adhesion and keratinocyte differentiation (Fig. 2a, b). According to the RNA-seq data, AJ molecules, such as E-cadherin and actin, were localized at intercellular junctions in areas of high cell density (Fig. 1c and 2c). E-cadherin and α-catenin form an AJ complex^9^, and through AJ development, α-catenin undergoes a conformational change due to intercellular forces, which is recognized by the α18 antibody^17^. α18 labeling was also pronounced in areas of high cell density (Fig. 2c). These data suggest that cells in regions of high cell density form AJs in response to intercellular forces.

**Figure 2.**
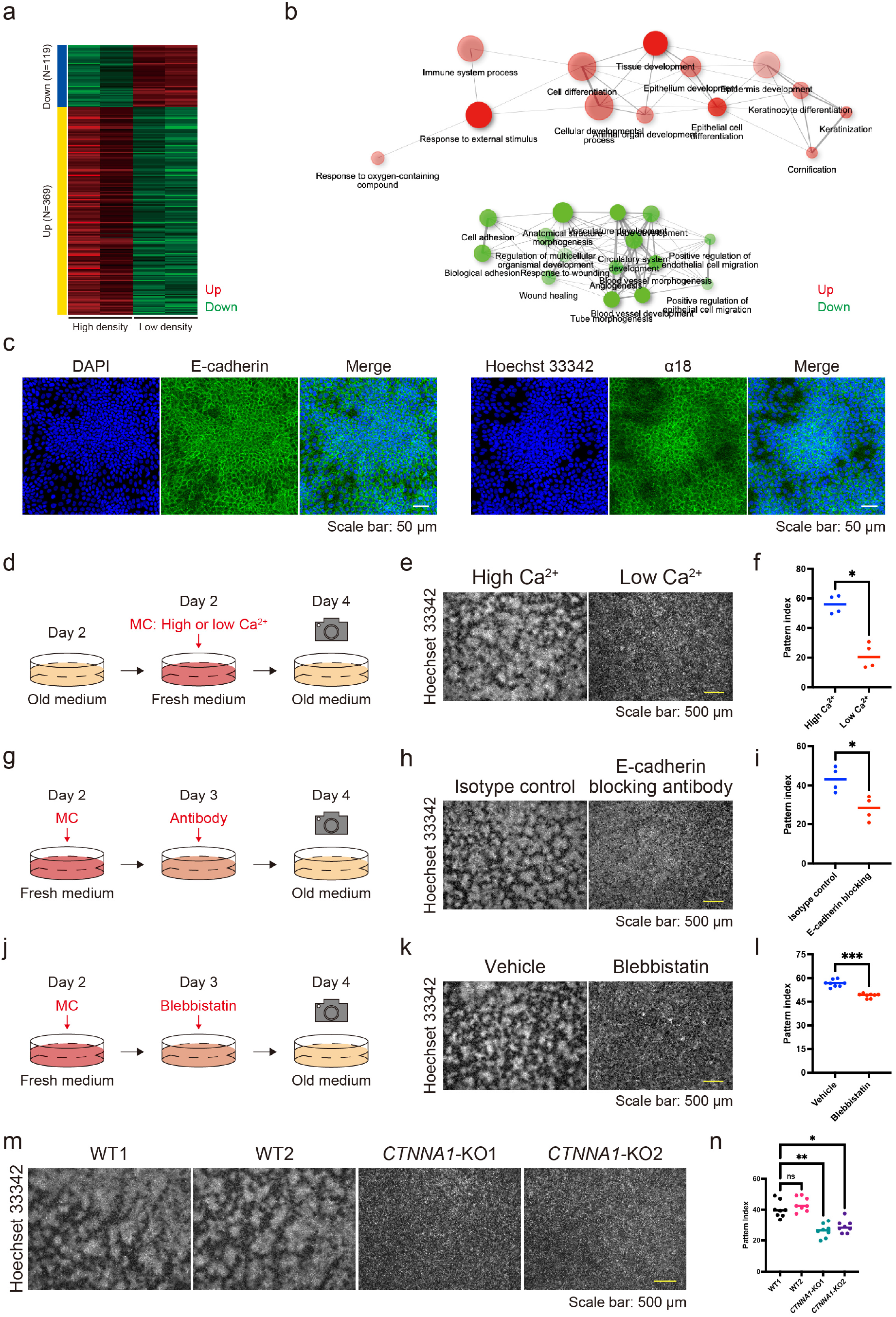
Adherens junctions regulate keratinocyte patterning. **a** Heatmap of differentially expressed genes (DEGs) identified by RNA sequencing (RNA-seq) comparing keratinocytes under a low- or high-density cell culture. **b** A network of gene ontology biological process terms enriched in DEGs. **c** Immunofluorescent images of keratinocytes on day 4. Nuclei are stained with 4′,6-diamidino-2-phenylindole (DAPI) and Hoechst 33342 (blue). Cell–cell adhesions are visualized with E-cadherin and α18 labeling (green). Scale bar: 50 µm. **d** Schematic diagram of experiments investigating patterns under high calcium (1.8 mM) and low calcium (0.06 mM) conditions. **e** Immunofluorescent images of keratinocytes under high and low calcium conditions. Nuclei are labeled with Hoechst 33342. Scale bar: 500 µm. **f** Pattern index in high and low calcium conditions. N = 4 for each group. **g** Schematic diagram of experiments investigating patterns with or without the E-cadherin-blocking antibody. **h** Immunofluorescent images of keratinocytes with or without the E-cadherin-blocking antibody. Nuclei are labeled with Hoechst 33342. Scale bar: 500 µm. **i** Pattern index with or without the E-cadherin-blocking antibody. N = 4 for each group. **j** Schematic diagram of experiments investigating patterns with or without blebbistatin, a non-muscle myosin II inhibitor. **k** Immunofluorescent images of keratinocytes with or without blebbistatin. Nuclei are labeled with Hoechst 33342. Scale bar: 500 µm. **l** Pattern index with or without blebbistatin. N = 8 for each group. **m** Immunofluorescent images of wild-type (WT) and *CTNNA1*-knockout (KO) keratinocytes. Nuclei are labeled with Hoechst 33342. Scale bar: 500 µm. **n** Pattern index of WT and *CTNNA1*-KO keratinocytes. N = 8 for each group. All data are presented as mean values. Data for **f**, **i**, and **l** were analyzed with two-tailed Mann–Whitney U tests. Data for **n** were analyzed with the Kruskal–Wallis test followed by Dunn’s multiple comparison test. ns, not significant. *, *P* < 0.05; **, *P* < 0.01; ***, *P* < 0.001.

To investigate the role of AJs in pattern development, we first compared cells under low- and high-calcium conditions because calcium is necessary to form AJs^18^. A high/low cell density pattern developed under high-calcium conditions but was not observed under low-calcium conditions (Fig. 2d–f). Similarly, treatment with SHE78-7 antibody, which blocks E-cadherin-mediated cell adhesion^19,20^, inhibited patterning (Fig. 2g–i). The force-dependent conformational change of α-catenin is induced by myosin-II^17^, and its inhibition by (-)-blebbistatin^21,22^ also disturbed pattern formation (Fig. 2j–l). *CTNNA1*-knockout (KO) HaCaT cell lines, in which α-catenin expression was nullified by gene editing (Supplementary Fig. 6), exhibited the same phenotype (Fig. 2m, n). These results demonstrate that AJs are essential for forming keratinocyte patterns.

### Cell–cell adhesion is sufficient for pattern development

We then asked whether cell–cell adhesion is sufficient for the formation of keratinocyte patterns and employed mathematical modeling to answer this question. We used a two-dimensional continuous model consisting of two variables: cell density and stress caused by adhesion. The main assumptions of the model were that the collective movement of cells is driven by the spatial imbalance of adhesion strength and that cell–cell adhesion increases with cell density. We performed simulations by varying the coefficient of adhesion strength as a control parameter. Each simulation began with spatially uniform density and stress. We observed that for sufficiently strong adhesion, the initial uniform distribution became unstable, and a spatial pattern of density emerged over time. By contrast, for weak adhesion, no coherent pattern appeared (Fig. 3; Supplementary Movie 5–7). These patterns were robust against noise strengths accounting for the random movements of individual cells (Supplementary Fig. 7). These results suggest that adequately strong cell–cell adhesion is sufficient for the emergence of density patterns.

**Figure 3.**
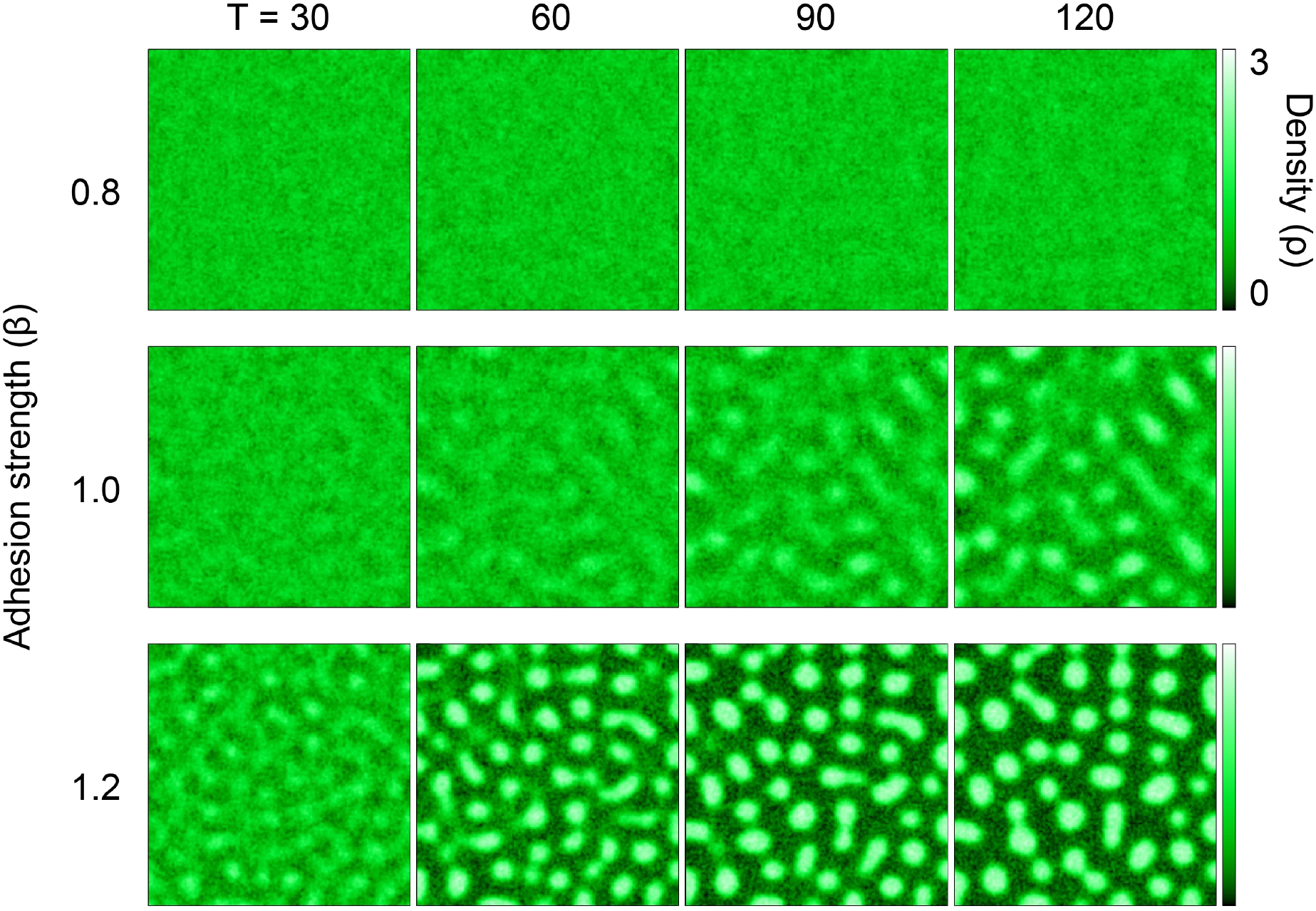
A mathematical model simulates a spatial density pattern using two variables: cell density and stress caused by adhesion. A two-dimensional cell density model predicted the emergence of spatial patterns of cell distribution, depending on adhesion strength. The color represents the local cell density ρ. Simulations for three different values of β (adhesion strength) are shown, with time moments T [a.u.] = 30, 60, 90, and 120.

### Keratinocyte patterns spatially dictate cell proliferation and differentiation

We further characterized keratinocyte differentiation through our experiments, as indicated by the RNA-seq data (Fig. 2b). Keratin 10 (KRT10)-positive differentiated cells were abundant and stratified in areas of high cell density (Fig. 4a, b). By contrast, phospho (Thr3)-monomethyl (Lys4) histone H3 (PH3)-positive proliferative cells were found in areas of low cell density (Fig. 4c). These data suggest that differentiation and proliferation are spatially regulated through pattern formation. Because AJs are essential for forming the high/low cell density pattern, we wondered whether they were also involved in regulating pattern-dependent differentiation and proliferation. *CTNNA1*-KO HaCaT cells failed to show pattern-dependent differentiation (Fig. 4d, e), suggesting that AJs involving α-catenin play a role in this process.

**Figure 4.**
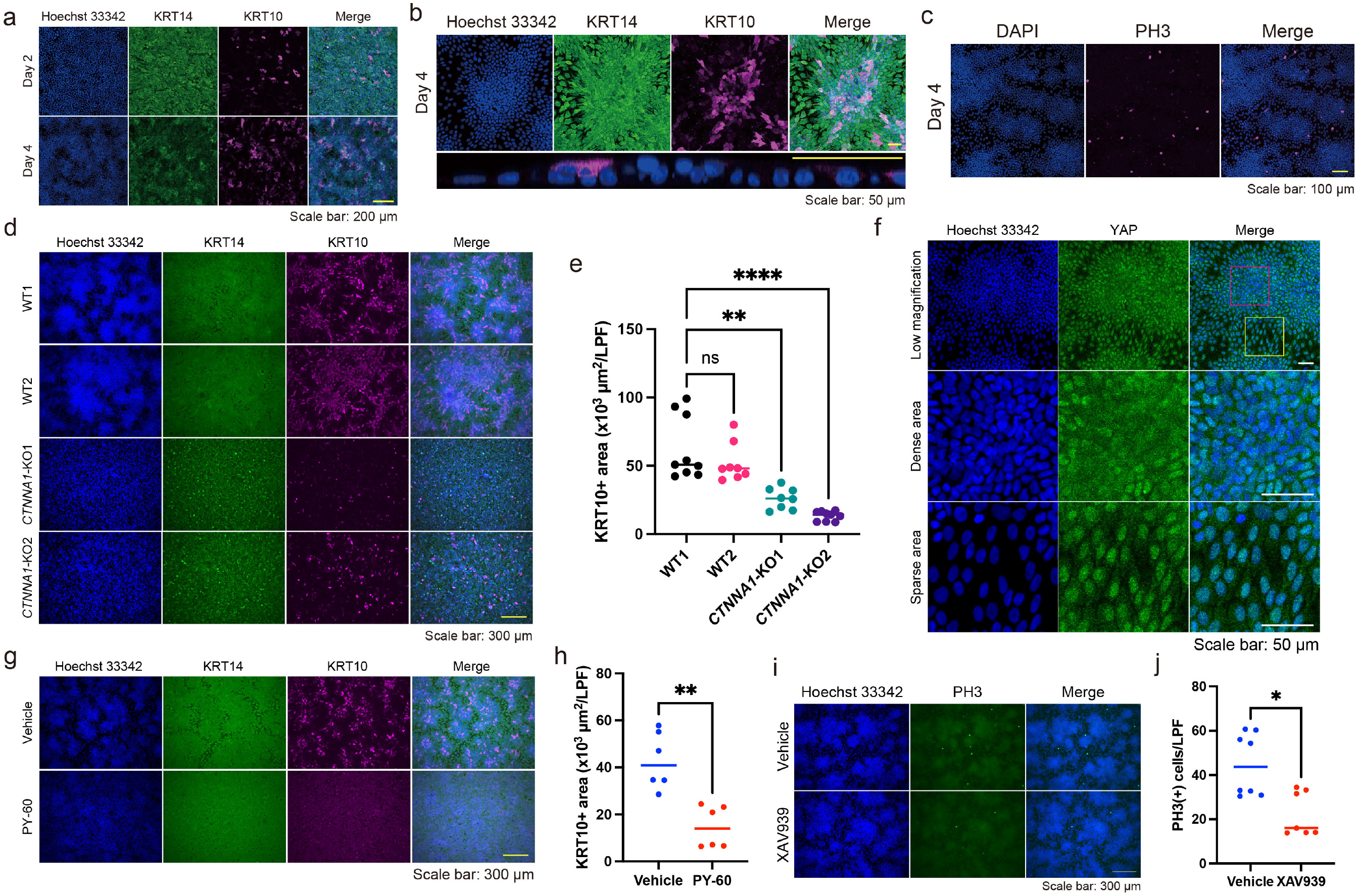
Patterning modulates differentiation and proliferation of keratinocytes. **a** Immunofluorescent images of keratinocytes on days 2 and 4. Hoechst 33342 (blue); Keratin (KRT) 14, a basal keratinocyte marker (green); KRT10, a differentiated keratinocyte marker (magenta). Scale bar: 200 µm. **b** The upper panels are immunofluorescent images of a high-density region on day 4. The lower panel is an orthogonal YZ image of a high-density region under high magnification. Hoechst 33342 (blue);KRT14 (green); KRT10 (magenta). Scale bar: 50 µm. **c** Immunofluorescent images of keratinocytes on day 4. DAPI (blue). PH3, a proliferation marker (magenta). Scale bar: 100 µm. **d** Immunofluorescent images of WT and *CTNNA1*-KO keratinocytes on day 4. Hoechst 33342 (blue); KRT14 (green); KRT10 (magenta). Scale bar: 300 µm. **e** KRT10-positive (KRT10+) areas per low power field (LPF) of WT and *CTNNA1*-KO keratinocytes per LPF. N = 9 for WT1 and *CTNNA1*-KO2. N = 8 for WT2 and *CTNNA1*-KO1. **f** Immunofluorescent images of keratinocytes on day 4. Hoechst 33342 (blue). Yes-associated protein (YAP), a cell density-sensitive molecule (green). The red and yellow squares indicate a dense area and a sparse area, respectively. Scale bar: 50 µm. **g** Immunofluorescent images of keratinocytes with or without PY-60, a YAP activator. Hoechst 33342 (blue); KRT14 (breen); KRT10 (magenta). Scale bar: 300 µm. **h** KRT10+ areas per LPF of immunofluorescent images for keratinocytes with or without PY-60. N = 6 for each group. **i** Immunofluorescent images of keratinocytes with or without XAV939, a tankyrase inhibitor. Hoechst 33342 (blue); PH3 (green). Scale bar: 300 µm. **j** Numbers of PH3-positive (PH3+) cells with or without XAV939. N = 8 for each group. All data are presented as mean values. Data for **e** were analyzed with the Kruskal–Wallis test followed by Dunn’s multiple comparison test. Data for **h** and **j** were analyzed with two-tailed Mann–Whitney U tests. ns, not significant. *, *P* < 0.05. **, *P* < 0.01. ****, *P* < 0.0001.

We then focused on the YAP pathway, as the coordinated pattern-dependent differentiation and proliferation seemed to be linked to cell density, and YAP, a “crowd control” molecule, uses α-catenin to sense cell density in keratinocytes^23^. The localization of YAP, which is regulated by cell density, determines cell fate; cytoplasmic YAP induces differentiation, and nuclear YAP promotes proliferation^23–26^. As expected, cytoplasmic YAP and nuclear YAP were found in areas of high and low cell density, respectively (Fig. 4f). The YAP activator PY-60 inhibited pattern-dependent differentiation (Fig. 4g, h). On the other hand, PY-60 also disrupted pattern formation (Supplementary Fig, 8a, b), which might have resulted from the impaired contact inhibition of proliferation^27^ or the dysregulation of AJs by YAP activation^28^. By contrast, YAP inhibition by a tankyrase inhibitor, XAV939, suppressed pattern-dependent proliferation (Fig. 4i, j), although XAV939 slightly modified pattern formation (Supplementary Fig. 8c, d). These results suggest that YAP modulates the differentiation and proliferation states of patterned keratinocytes.

### Serum starvation contributes to epidermal stratification

Finally, we asked whether the keratinocyte patterns induced by serum starvation impacted the three-dimensional (3D) organization of the epidermis, as cells in areas of high cell density within the pattern were predisposed to differentiation. The 3D culture of HaCaT cells revealed that the epidermis was thicker in serum-starved conditions than in serum-rich conditions, in which pattern formation was impaired (Fig. 5a, b; Supplementary Fig. 9a, b). In addition, *CTNNA1*-KO HaCaT cells failed to form a stratified epithelium (Fig. 5c).

**Figure 5.**
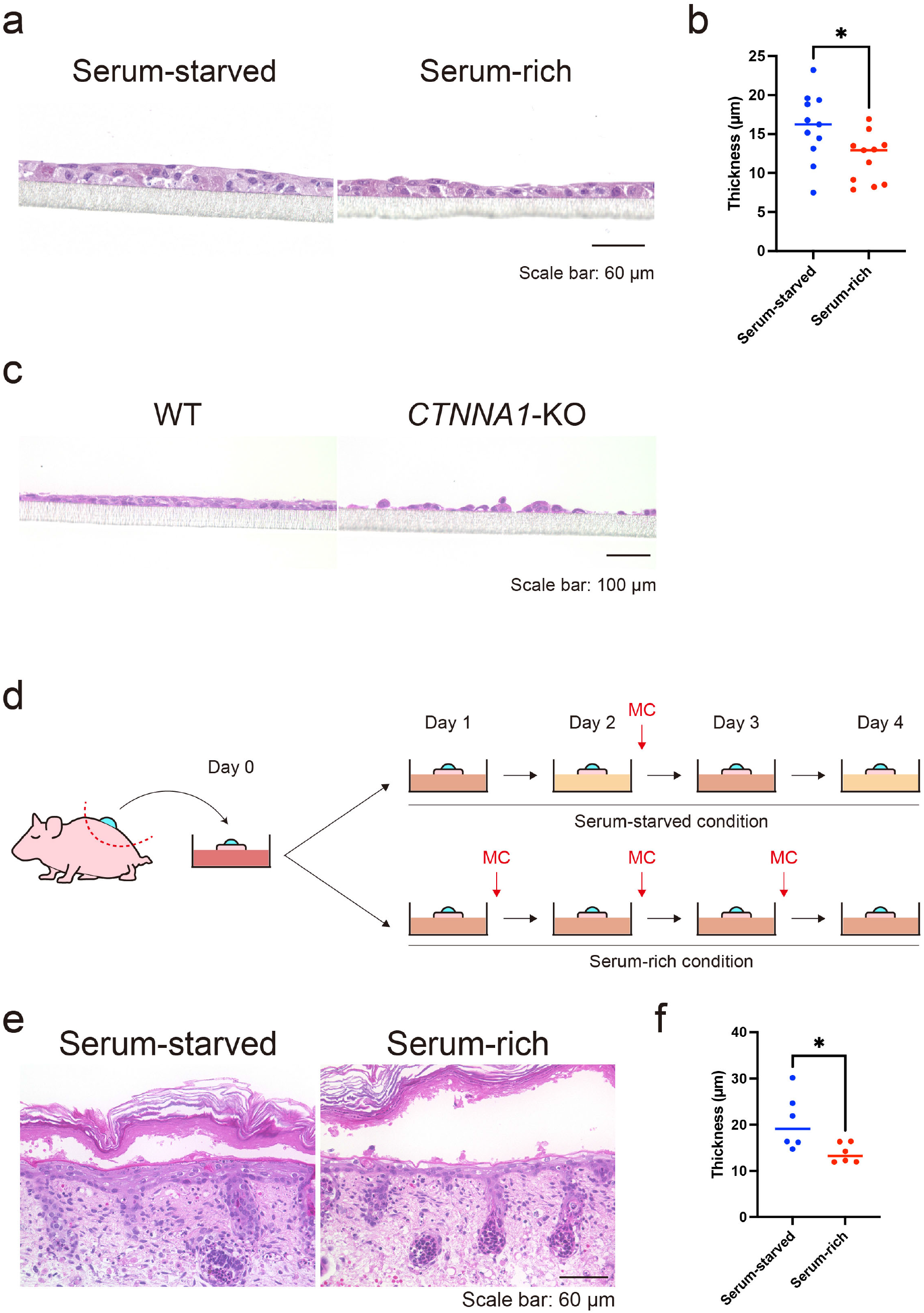
Three-dimensional *in vitro* and *ex vivo* epidermis cultures under serum-starved and serum-rich conditions. **a** Hematoxylin and eosin (H&E) staining of 3D epidermis cultures in serum-starved and serum-rich conditions. Scale bar: 60 µm. **b** Quantification of epidermal thickness in the 3D culture under serum-starved and serum-rich conditions. N = 11 for each group. **c** H&E staining of WT and *CTNNA1*-KO 3D epidermis cultures. Scale bar: 100 µm. **d** Schematic diagram of *ex vivo* culture experiment using P1 neonate back skin with suction blister wound under serum-starved and serum-rich conditions. **e** H&E staining of *ex vivo* cultured tissue. Scale bar: 60 µm. **f** Quantified re-stratified epidermal thickness in the *ex vivo* culture. N = 6 for each group. All data are presented as mean values and were analyzed with two-tailed Mann–Whitney U tests. *, *P* < 0.05.

**Figure 6.**
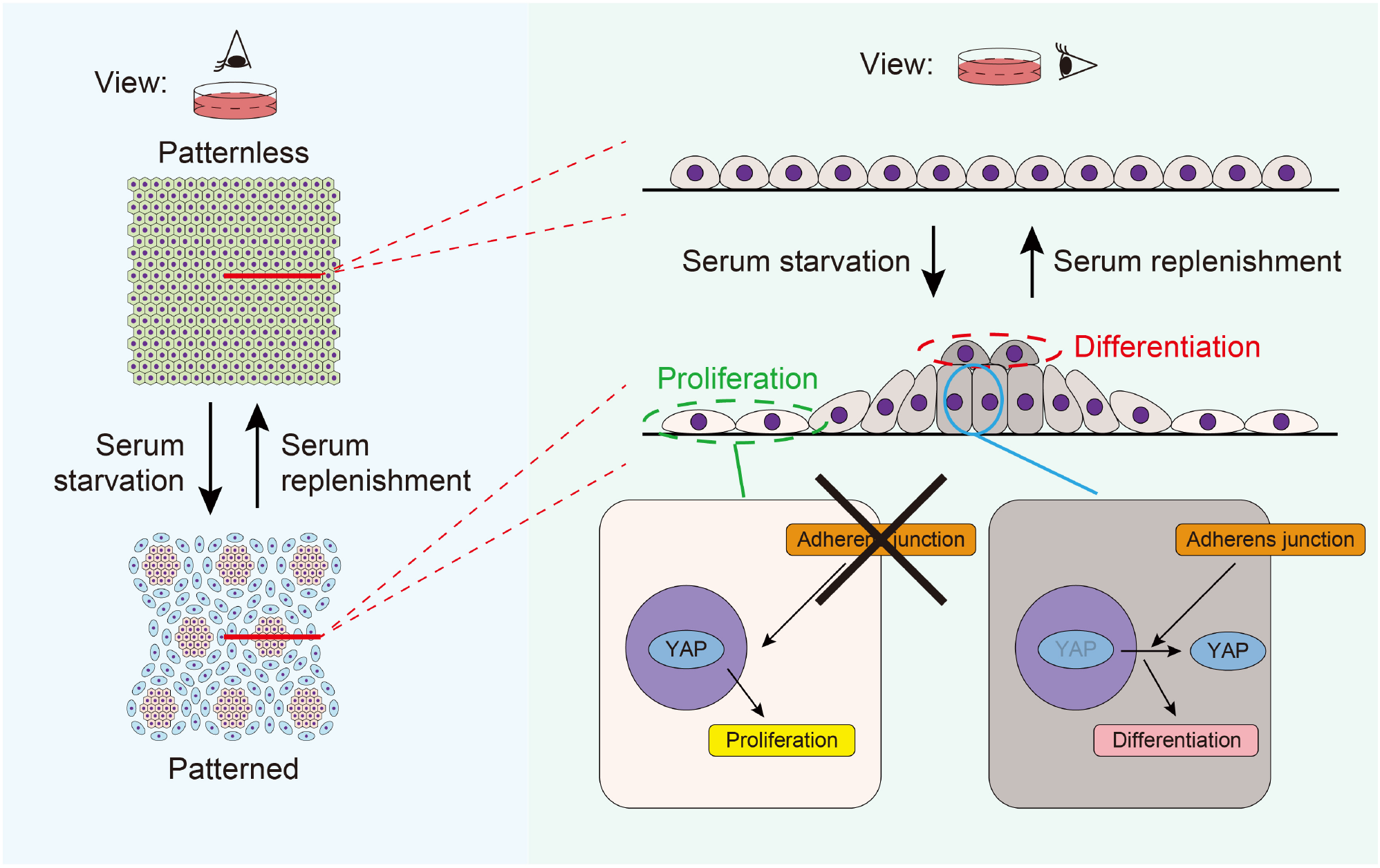
Graphical summary of keratinocyte patterning. Serum starvation induces keratinocyte patterns, comprising areas of high and low cell density. This pattern formation is reversible upon serum replenishment. Adherens junctions (AJs) regulate this patterning. The low- and high-density pattern spatially dictates keratinocyte differentiation and proliferation through YAP, which senses cell density.

We further evaluated whether the serum-starved condition could affect epidermal stratification *ex vivo*. We employed a wound-healing mouse model with a suction blister to observe re-stratification after wounding^29^. A sample of mouse back skin with a suction blister wound was cut out and cultured in serum-starved and serum-rich conditions (Fig. 5d). Re-epithelialization was completed on day 1, and re-stratification began. On day 4, the re-stratified epidermis cultured in serum-starved conditions was thicker than that cultured in serum-rich conditions (Fig. 5e, f). These data imply that appropriate serum starvation facilitates epidermal stratification through cell–cell adhesion and the subsequent patterning of the cells.

## Discussion

Our study presents a novel and robust model of cell–cell adhesion-induced patterning (CAIP). CAIP is mediated by AJs and spatially regulates the differentiation and proliferation of epithelial cells.

During skin development, such as placode formation, the periodic expression of signaling molecules is thought to arise in the early stages of development, dictate cell behaviors, and orchestrate follicular patterning in the skin^30–32^. However, recent studies have revealed that periodic follicle patterning is triggered by mechanical rather than molecular events^2,33^. In this scenario, follicular patterns can arise from mechanical instability caused by fibroblast contraction. Then, self-organized fibroblast aggregation through contractility-driven cellular pulling triggers the mechanosensitive activation of β-catenin in neighboring keratinocytes, activating the follicle gene expression program. These studies indicate that cellular contraction mechanics in the mesenchyme could organize epithelial patterning and subsequent cell fate decisions *in vivo*. In contrast to the follicle placodes, fingerprint ridge formation, which involves sweat gland development^1^, is facilitated by signaling molecules, including EDAR, WNT, and BMP, and does not recruit mesenchymal cells. The concept of CAIP is distinct because epithelial cell–cell adhesion is essential and sufficient for the patterning. Research to determine whether CAIP is involved in other *in vivo* epithelial patterning is warranted.

CAIP also imitates the periodic buckling of the human epidermis, in which epidermal stem cells with high β1 integrin expression are localized in the areas facing dermal protrusion and transient amplifying cells are present in the epidermal rete ridge structures^34,35^. Keratinocytes cultured on undulating elastomer substrates resembling rete ridge structures display a similar stem cell distribution, in which cells with high β1 integrin expression cluster at the tips of the topographies^36,37^. The disruption of cell–cell adhesions by Rho kinase inhibitors impairs confluent sheet formation on undulating substrates. However, the stem cell clusters on undulating substrates strongly express E-cadherin and F-actin, in contrast to the high E-cadherin expression in the differentiation region of CAIP. This discrepancy might be explained by differences in the experimental designs.

Serum starvation induces quiescence *in vitro*^38^ and has been used for cultured keratinocytes^39–41^, fibroblasts^42,43^, and other cell types^44–46^. By contrast, serum stimulation enhances keratinocyte migration^39,47,48^. Hence, CAIP in serum-starved conditions might reflect the resting state of the epidermis. However, in our study, CAIP and serum starvation also supported epithelial stratification, which could simulate epidermal morphogenesis or the final stage of wound healing in the epidermis. These data indicate that CAIP modulation might help finalize wound closure by enhancing epidermal stratification once the wound gap is filled with one layer of epithelial cells. In line with this, evidence indicates that hyperhydration or excessive extracellular fluid delays wound closure in clinical settings^49,50^. Theoretically, the serum-starved culture, which reduces the MC frequency, could be beneficial to other epidermis or skin organoid cultures to obtain thicker epidermal sheets efficiently and economically. Our findings may also have applications in regenerative medicine, such as the preparation of epidermal grafts.

In conclusion, our study uncovered the pivotal role of cell–cell adhesion in driving epithelial cell patterning. We propose that the molecules involved in CAIP are suitable target candidates to facilitate wound healing.

## Methods

### Cells

HaCaT cells^14^ were originally obtained from Dr. Norbert Fussenig (German Cancer Research Center, Heidelberg, Germany), and the cell identity was confirmed with the Cell Culture STR profile (Biologica, Nagoya, Japan). Single-cell-cloned HaCaT cells with stable Cas9 expression were established previously^15^. Briefly, HaCaT cells were transfected with pLenti-EF1a-Cas9-Puro lentiviral particles (Applied Biological Materials, Richmond, Canada) and selected with 1 µg/mL puromycin (Thermo Fischer Scientific, Waltham, Massachusetts, US). Thereafter, 50 cells were seeded into a 10 cm dish, and the single-cell clone was obtained with Scienceware cloning discs (Sigma-Aldrich, Darmstadt, Germany) dipped in trypsin (Wako, Osaka, Japan). To establish *CTNNA1* HaCaT cells, crRNA was acquired from predesigned Alt-R CRISPR-Cas9 guide RNA (Integrated DNA Technologies, Coralville, Iowa, US); the crRNA contained the TGAAGCGAGGCAACATGGTT sequence with the CGG PAM sequence targeting exon 4 of the *CTNNA1* genomic sequence (NG_047029.1). For transfection, HaCaT cells stably expressing Cas9 were seeded onto a 24-well plate with a concentration of 300 × 10^5^ cells/well. One day after seeding, HaCaT cells were transfected with a 10 nM duplex of crRNA and Alt-R CRISPR-Cas9 tracrRNA tagged with ATTO 550 (Integrated DNA Technologies) using Lipofectamine RNAiMAX Transfection Reagent (Thermo Fisher Scientific) according to the manufacturer’s instructions. Thereafter, the single-cell clone was obtained by seeding onto a 96-well plate. The truncations on the *CTNNA1* gene caused by CRISPR/Cas9 were confirmed by Sanger sequencing, and the knockout of the *CTNNA1* gene was examined by Western blot. All cell stocks were routinely tested for mycoplasma contamination, and all tests were negative.

### Cell culture

HaCaT cells were maintained in Dulbecco’s modified eagle medium (DMEM) with 4.5 g/L glucose and L-glutamine (Nacalai Tesque, Kyoto, Japan) supplemented with 10% FBS (HyClone, Marlborough, Massachusetts, US) and 1x antibiotic–antimycotic mixed stock solution (Nacalai Tesque) at 37 °C in humidified air with 5% CO2. For the live imaging with confocal microscopy, 4.5 g/L glucose DMEM without phenol red (Nacalai Tesque) supplemented with L-glutamine, 10% FBS, and 1x antibiotic–antimycotic mixed stock solution was used for cell culture. Low-calcium-condition experiments utilized calcium-free 4.5 g/L glucose DMEM without L-glutamine (Nacalai Tesque) or calcium- and phenol red-free 4.5 g/L glucose DMEM without L-glutamine supplemented with 10% calcium-free FBS. To prepare calcium-free FBS, 5 g Chelex 100 (Bio-Rad, Hercules, California, US) was added to 100 mL FBS with shaking for 1 hour at room temperature, followed by filtration with a 0.22 µm Stericup-GP Express Plus PES (Millipore, Darmstadt, Germany).

### Pattern formation of cultured keratinocytes

HaCaT cells were seeded at 5.0 × 10^5^ cells/mL onto µ-Slide 8 Well Chambers (ibidi, Martinsried, Germany) using 250 µL of suspension per well or onto a µ-Dish 35 mm dish (ibidi, Martinsried, Germany) using 2 mL of suspension per dish. The culture medium was replenished every 2 days with equivalent volumes unless otherwise specified. The keratinocyte pattern was observed 4 days after seeding.

### Inhibitor treatments

HaCaT cells were treated with the E-cadherin-blocking antibody (SHE78-7, Takara, Shiga, Japan), PY-60 (Axon Medchem, Groningen, Netherlands), or XAV939 (Fujifilm, Tokyo, Japan) at a concentration of 30 µg/mL when the culture medium was replenished 2 days after seeding. The cells were observed 2 days after treatment. Blebbistatin (Cayman Chemical, Ann Arbor, Michigan, US) was used at a concentration of 12.5 µM 3 days after seeding. The cells were evaluated 4 days after seeding.

### Three-dimensional epidermal culture

ThinCert 12-well cell culture inserts with 0.4 µm pores (Greiner Bio-One, Kremsmünster, Austria) were placed on Falcon 12-well cell culture plates (Corning, Corning, New York, US) and precoated with CTS CELLStart Substrate (Thermo Fischer Scientific) in a 1:50 dilution of Dulbecco’s phosphate-buffered saline (PBS) with MgCl2 and CaCl2 (Sigma-Aldrich) overnight. Subsequently, 1 mL of culture media was added into the well, and 1 mL of HaCaT cell suspension at a concentration of 2.5 × 10^5^ cells/mL was seeded onto the inserts. In the serum-starved condition, the culture medium for the insert and the well was replenished two days after seeding. In the serum-rich condition, the medium was replenished every day. For placing the inserts under an air-liquid interface culture, Extra Thick Blot Filter Paper (Bio-Rad) was cut to 2.8 cm × 2.8 cm with two 12 mm holes. The filter paper was placed into a well of Falcon 6-well deep well plates (Corning), which was filled with 11.5 mL of the cell culture medium. Four days after seeding, the culture medium within the insert was removed in both conditions, and the inserts were placed on the filter papers. The culture medium in the well was replenished every 3 days for the serum-starved condition and every day for the serum-rich condition for 14 days after seeding. Samples were collected by excising the membranes from the inserts and were fixed with formalin.

### *Ex vivo* skin culture

Suction blisters were generated on neonatal C57BL/6 murine dorsal skin (P1) using a syringe and connector tubes^29^. The back skin with the suction blister was excised and cultured with DMEM with 4.5 g/L glucose and L-glutamine supplemented with 10% FBS and 1x antibiotic–antimycotic mixed stock solution. In the serum-starved condition, the culture medium was changed on the second day of cultivation. In the serum-rich condition, the medium was replenished every day. Samples were fixed with formalin on the fourth day of cultivation.

### Live cell imaging

HaCaT cells at a concentration of 5.0 × 10^5^ cells/mL were seeded onto µ-Slide 8 well chambers (250 µL per well) and 3.5 cm plastic dishes (2 mL per dish). HaCaT cells were cultured as indicated in the figures, and live cell imaging was carried out with a BZ-9000 or BZ-X800 microscope (Keyence, Osaka, Japan) equipped with an incubation chamber maintained at 37 °C and 5% CO2.

### Autoperfusion system

HaCaT cells with a concentration of 5.0 × 10^5^ cells/mL were seeded onto 3.5 cm plastic dishes (2 mL per dish). Two days after seeding, the culture medium was replenished, and culture dishes were connected to the autoperfusion system. The autoperfusion system consisted of a micro tube pump system (iCOMES Lab, Iwate, Japan) and 50 mL conical tubes that supplied the fresh culture medium and discarded the used culture medium. The perfusion rates were 2 mL/2 days for the low perfusion group and 6 mL/2 days for the high perfusion group, simulating serum-starved and serum-rich conditions, respectively. Samples were fixed with 4% paraformaldehyde at room temperature for 10 minutes.

### Mathematical model

Consider a two-dimensional continuous model with cell density *ρ*(*x*, *y*, *t*) and negative pressure *σ*(*x*, *y*, *t*) (the diagonal component of the stress tensor) created by the accumulation of AJ complexes. The time evolution of these variables is governed by:

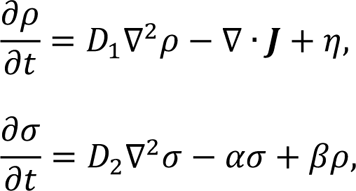

where 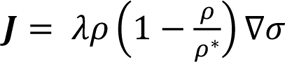 represents the active cell flow due to force balance, which vanishes when the local cell density reaches the carrying capacity *ρ*^∗^. We set *ρ*^∗^ = 3, allowing local squeezing and overlapping of cells of up to 300%. The variable *η* accounts for the random motion of cells. To perform simulations, the two-dimensional space of size [*L*, *L*] was discretized into a 200 × 200 grid. To preserve the total cell volume, the random force at grid (*i*, *j*) was chosen as *η*(*x_i_*, *y_j_*, *t*) = *N*(*ξ*_*i*+1,*j*_+*ξ*_*i*−1,*j*_+*ξ*_*i,j*+1,*j*_*ξ*_*i,j*−1,*j*_4*ξ*_*i,i*_), where *N* is the noise strength and *ξ_i,j_* is a Gaussian random variable with zero mean and unit variance. The equation for stress (*σ*) is a natural extension of a one-dimensional model derived by Recho et al.^51^, in which the off-diagonal component of the stress tensor is neglected. The stress *σ* is assumed to be proportional to the concentration of AJs, which accumulates with the rate *β* as the cell density increases, consistent with the observation that regions of high cell density form AJs in response to intercellular forces. It also decays at the rate *α*. The resulting model is a variant of the Keller–Segel system^52^. A saturation term in the active flow in our model ensures that the density remains finite, as it diverges in the original Keller–Segel system.

*β* was chosen as a control parameter. For simulations, the parameters wer set as follows: *L* = 200, *D*_1_ = 0.5, *D*_2_ = 5.0, *α* = 1.0, *λ* = 1.0, and *N* = 0.5 (for Figure 3) or *N* ∈ {0.1, 0.5, 1.0} (for Supplementary Figure 7). For all simulations, a spatially uniform initial condition *ρ*(*x*, *y*, *t*) = 1 and *σ*(*x*, *y*, *t*) = 0 was chosen (which means that cells are confluent in the system and no AJs are expressed) with the flux-free boundary condition.

### Animals

C57BL/6 mice were purchased from Clea Japan (Tokyo, Japan). The institutional review board of the Hokkaido University Faculty of Medicine and Graduate School of Medicine approved all animal experiments in this study.

### Histology

For the epidermal thickness analysis, membrane samples from inserts or *ex vivo* cultured back skins were embedded in paraffin after dehydration. Thereafter, sectioned paraffin samples were deparaffinized and stained with hematoxylin and eosin (H&E). Images were taken with a BZ-9000 microscope, and the thickness of the epidermis was analyzed by ImageJ.

### Immunofluorescent staining

The cells were cultured on µ-Slide 8 well chambers or µ-Dish 35 mm dishes (ibidi, Martinsried, Germany). The cells were washed with PBS (Nacalai Tesque) and fixed with 4% paraformaldehyde at room temperature for 10 minutes. Cells were permeabilized with 0.1% or 0.5% Triton X-100 in PBS for 20 minutes at room temperature, followed by blocking with 3% bovine serum albumin in PBS for 30 minutes. Subsequently, cells were incubated for 1–2 hours at room temperature with the following primary antibodies: anti-E-cadherin antibody (Cell Signaling Technology, Danvers, Massachusetts, US, Cat# 3195, RRID:AB_2291471, 1:100 dilution), anti-α18 antibody^17^, anti-PH3 antibody (Millipore, Cat# 07-554, RRID:AB_11210699, 1:250 dilution), anti-K14 antibody (Thermo Fischer Scientific, Cat# MA5-11599, RRID:AB_10982092), anti-K10 antibody (BioLegend, San Diego, California, US, Cat# 905404, RRID:AB_2616955), and anti-YAP antibody (Cell Signaling Technology, Cat# 14710, RRID:AB_2798583). Secondary antibodies, namely goat anti-mouse IgG Alexa Fluor 546 (Thermo Fischer Scientific, Cat# A11003, RRID:AB_2534071, 1:1,000 dilution) and goat anti-rabbit IgG Alexa Fluor 488 (Thermo Fischer Scientific, A21206, RRID:AB_2535792, 1:1,000 dilution), were incubated for 1 hour at room temperature. Cells were washed in PBS 3 times for 5 minutes. Nuclei were stained with 4’,6-diamidino-2-phenylindole (Thermo Fischer Scientific) or Hoechst 33342 (Dojindo, Kumamoto, Japan) at a concentration of 0.5 µg/mL or 5 µg/mL for 1 hour at room temperature. Photo images were captured using BZ-9000 (Keyence), FV-1000 (Olympus, Tokyo, Japan), or LSM 710 (Zeiss, Oberkochen, Germany) imaging systems. Phalloidin-iFluor 555 Reagent (Abcam, Cambridge, UK, Cat# ab176756, 1:500 dilution) was used with Hoechst 33342 for actin staining at the time of primary antibody incubation. After washing with PBS, cells were observed with an LSM 710 microscope. In the autoperfusion experiment, cells were washed with PBS and stained with 0.5 µg/mL Hoechst 33342 at room temperature for 20 min. After PBS washes, images were taken using a BZ-9000 microscope. To observe the pattern of the 3D epidermis culture, the fixed membranes were washed with PBS and stained with 0.5 µg/mL Hoechst 33342 at room temperature for 1 hour. Thereafter, inserts were washed with PBS and mounted in Fluoromount-G (Thermo Fischer Scientific). Images were obtained with an LSM 710 microscope.

### Western blot

Wild-type (WT) or *CTNNA1*-KO HaCaT cells at a concentration of 1.0 × 10^6^ cells/well were seeded onto 6-well plastic plates. Two days after seeding, we collected cell lysates using a lysis buffer containing 1% Nonidet P-40 (Nacalai Tesque), 25 mM Tris-HCl (pH 7.4), 100 mM NaCl, 10 mM ethylenediaminetetraacetic acid, and a 1:100 dilution protease inhibitor cocktail (P8340, Sigma-Aldrich) on ice for 30 minutes with shaking. The whole cell lysates were centrifuged at 13,000 rpm at 4 °C for 20 minutes. Cell lysate supernatants were denatured with a 5× loading buffer (0.25 M Tris-HCl; 8% sodium dodecyl sulfate; 30% glycerol; 0.02% bromophenol blue; 0.3 M β-mercaptoethanol; pH 6.8). Samples were subjected to SDS-PAGE using NuPAGE 4 to 12%, Bis-Tris, 1.0–1.5 mm, Mini Protein Gels. Proteins separated by SDS-PAGE were electrophoretically transferred onto PVDF transfer membranes (Bio-Rad). The membranes were blocked with 2% skim milk and incubated with rabbit ant-α-catenin antibodies (Thermo Fischer Scientific, Cat# 71-1200, RRID:AB_2533974, 1:200 dilution) or rabbit anti-β-tubulin antibodies (Abcam, Cat# ab6046, RRID:AB_2210370, 1:5000 dilution) in 2% skim milk at room temperature for 1 hour. After washing with Tris-buffered saline, the membranes were incubated with a dilution of 1:5000 peroxidase-conjugated anti-rabbit IgG antibody (Jackson ImmunoResearch, West Grove, Pennsylvania, US, AB_10015282) in 2% skim milk at room temperature for 1 hour. Signals were visualized by Clarity Western ECL Substrate (Bio-Rad) and detected with an ImageQuant LAS 4000 mini camera system (Fujifilm).

### RNA-seq

HaCaT cells were seeded into 6 cm plastic dishes at a concentration of 2.5 × 10^5^/mL or 7.5 × 10^5^/mL. After overnight incubation at 37 °C in humidified 5% CO2, RNA was extracted using the FastGene RNA Premium Kit (Nippon Genetics, Tokyo, Japan) according to the manufacturer’s instructions. Library preparation and sequencing were performed by Novogene (Beijing, China). Briefly, mRNA was enriched using oligo(dT) beads. The mRNA was fragmented randomly by adding a fragmentation buffer, then the cDNA was synthesized using an mRNA template and a random hexamer primer, after which a custom second-strand synthesis buffer (Illumina, San Diego, California, US), dNTPs, RNase H, and DNA polymerase I were added to initiate second-strand synthesis. After a series of terminal repair, A-tailing, and sequencing adaptor ligation, the double-stranded cDNA library was completed through size selection and PCR enrichment. After the quality control of the libraries, the libraries were sequenced by the NovaSeq 6000 system in PE150 mode. Reads were then mapped to hg38 using STAR (v.2.7.3)^53^. Gene expression levels were quantified using RSEM (v.1.3.1)^54^. Read counts were analyzed through integrated differential expression and pathway analysis (iDEP9.51)^55^. Genes with low-level expression (less than 0.5 counts per million in all samples) were removed from the analysis. Genes that were differentially expressed between low-density and high-density conditions were identified with DESeq2 using a threshold of false discovery rate of <0.1 and a fold-change of >2^16^. The upregulated and downregulated genes were subjected to enrichment analysis.

### Calculation of Pattern Index

Grayscale versions for immunofluorescent images of keratinocytes, in which the nuclei were labeled with Hoechst 33342, were used to evaluate the inhomogeneity of cell density. The image was binarized with the threshold value determined from its brightness value histogram (i.e., the threshold value was defined as the brightness that had the maximum count and was greater than 0.6 times the average brightness of the total image). Subsequently, the binarized image was blurred using a linear filter within a circle with a radius *R*. Here, *R* was set to 4 pixels, corresponding to approximately 10 µm, which was nearly the same as the distance between neighboring cells. It should be noted that the blurred image was almost uniform if the cells were uniformly distributed, but it had different values if the cell density changed. Thus, the standard of the blurred images was calculated and adopted as the pattern index, in which a greater value indicates a more spatially inhomogeneous distribution of cells.

### Statistical analysis

Statistical analyses were performed with GraphPad Prism 9 (GraphPad Software). *P*-values were determined with two-tailed Mann–Whitney U tests or Kruskal–Wallis tests followed by Dunn’s multiple comparison test. *P*-values of <0.05 were considered statistically significant.

## Supporting information

Description of additional supplementary files

## Acknowledgments

We thank Nami Ikeshita and Mika Tanabe for their technical assistance. We also thank Professor Akira Nagafuchi for providing α18 antibodies. This work was funded by JSPS KAKENHI Grant Number 23K15277, the Tokyo Biochemical Research Foundation, the Nakatomi Foundation to YM, JSPS KAKENHI Grant Number JP22K03428 to YK, JST CREST Grant Number JPMJCR1926, JSPS KAKENHI Grant Number JP23H04936 to MN, JSPS KAKENHI Grant Number 23H02928, the Akiyama Life Science Foundation, and the Mochida Memorial Foundation for Medical and Pharmaceutical Research to KN.

## Author contributions

Y.M. performed the experiments, analyzed the data, interpreted the results, and wrote the manuscript. Y.K. developed the mathematical model, interpreted the results, and wrote the manuscript. H.K. calculated the pattern index and wrote the manuscript. T.S., T. N., S.I., and S.M. performed the experiments. J.K. designed the experiments. M.N. interpreted the results and supervised the study. W.N. performed the experiments, analyzed the data, and interpreted the results. H.U. supervised the study. K.N. interpreted the results, wrote the manuscript, and supervised the study.

## References

1. Glover, J. D. et al. The developmental basis of fingerprint pattern formation and variation. Cell 186, 940–956.e20 (2023).

2. Shyer, A. E. et al. Emergent cellular self-organization and mechanosensation initiate follicle pattern in the avian skin. Science 357, 811–815 (2017).

3. Gomez, C. et al. The Interfollicular Epidermis of Adult Mouse Tail Comprises Two Distinct Cell Lineages that Are Differentially Regulated by Wnt, Edaradd, and Lrig1. Stem Cell Rep. 1, 19–27 (2013).

4. Sada, A. et al. Defining the cellular lineage hierarchy in the interfollicular epidermis of adult skin. Nat. Cell Biol. 18, 619–631 (2016).

5. Wang, Y. et al. Collagen XVII deficiency alters epidermal patterning. Lab Invest 102, 581–588 (2022).

6. Green, H. & Thomas, J. Pattern Formation by Cultured Human Epidermal Cells: Development of Curved Ridges Resembling Dermatoglyphs. Science 200, 1385–1388 (1978).

7. Jones, P. H., Harper, S. & Watt, F. M. Stem cell patterning and fate in human epidermis. Cell 80, 83–93 (1995).

8. Klein, A. M., Nikolaidou-Neokosmidou, V., Doupé, D. P., Jones, P. H. & Simons, B. D. Patterning as a signature of human epidermal stem cell regulation. J Roy Soc Interface 8, 1815–1824 (2011).

9. Simpson, C. L., Patel, D. M. & Green, K. J. Deconstructing the skin: cytoarchitectural determinants of epidermal morphogenesis. Nat Rev Mol Cell Bio 12, 565–580 (2011).

10. Larue, L., Ohsugi, M., Hirchenhain, J. & Kemler, R. E-cadherin null mutant embryos fail to form a trophectoderm epithelium. Proc. Natl. Acad. Sci. 91, 8263–8267 (1994).

11. Huelsken, J. et al. Requirement for β-Catenin in Anterior-Posterior Axis Formation in Mice. J. Cell Biol. 148, 567–578 (2000).

12. Torres, M. et al. An α-E-catenin gene trap mutation defines its function in preimplantation development. Proc. Natl. Acad. Sci. 94, 901–906 (1997).

13. Davis, M. A. & Reynolds, A. B. Blocked Acinar Development, E-Cadherin Reduction, and Intraepithelial Neoplasia upon Ablation of p120-Catenin in the Mouse Salivary Gland. Dev. Cell 10, 21–31 (2006).

14. Boukamp, P. et al. Normal keratinization in a spontaneously immortalized aneuploid human keratinocyte cell line. J Cell Biology 106, 761–771 (1988).

15. Mai, S. et al. Native Autoantigen Complex Detects Pemphigoid Autoantibodies. JID Innovations 3, 100193 (2023).

16. Love, M. I., Huber, W. & Anders, S. Moderated estimation of fold change and dispersion for RNA-seq data with DESeq2. Genome Biol 15, 550 (2014).

17. Yonemura, S., Wada, Y., Watanabe, T., Nagafuchi, A. & Shibata, M. α-Catenin as a tension transducer that induces adherens junction development. Nat Cell Biol 12, 533–542 (2010).

18. Vasioukhin, V., Bauer, C., Yin, M. & Fuchs, E. Directed Actin Polymerization Is the Driving Force for Epithelial Cell–Cell Adhesion. Cell 100, 209–219 (2000).

19. Shirayoshi, Y., Nose, A., Iwasaki, K. & Takeichi, M. N-Linked Oligosaccharides Are Not Involved in the Function of a Cell-Cell Binding Glycoprotein E-Cadherin. Cell Struct Funct 11, 245– 252 (1986).

20. Watabe, M., Nagafuchi, A., Tsukita, S. & Takeichi, M. Induction of polarized cell-cell association and retardation of growth by activation of the E-cadherin-catenin adhesion system in a dispersed carcinoma line. J Cell Biology 127, 247–256 (1994).

21. Kovács, M., Tóth, J., Hetényi, C., Málnási-Csizmadia, A. & Sellers, J. R. Mechanism of Blebbistatin Inhibition of Myosin II*. J Biol Chem 279, 35557–35563 (2004).

22. Limouze, J., Straight, A. F., Mitchison, T. & Sellers, J. R. Specificity of blebbistatin, an inhibitor of myosin II. J Muscle Res Cell Motil 25, 337–341 (2004).

23. Schlegelmilch, K. et al. Yap1 Acts Downstream of α-Catenin to Control Epidermal Proliferation. Cell 144, 782–795 (2011).

24. Zhang, H., Pasolli, H. A. & Fuchs, E. Yes-associated protein (YAP) transcriptional coactivator functions in balancing growth and differentiation in skin. Proc National Acad Sci 108, 2270–2275 (2011).

25. Zhao, B. et al. Inactivation of YAP oncoprotein by the Hippo pathway is involved in cell contact inhibition and tissue growth control. Gene Dev 21, 2747–2761 (2007).

26. Totaro, A. et al. YAP/TAZ link cell mechanics to Notch signalling to control epidermal stem cell fate. Nat Commun 8, 15206 (2017).

27. Shalhout, S. Z. et al. YAP-dependent proliferation by a small molecule targeting annexin A2. Nat. Chem. Biol. 17, 767–775 (2021).

28. Neto, F. et al. YAP and TAZ regulate adherens junction dynamics and endothelial cell distribution during vascular development. Elife 7, e31037 (2018).

29. Fujimura, Y. et al. Hair follicle stem cell progeny heal blisters while pausing skin development. EMBO Rep. 22, e50882 (2021).

30. Widelitz, R. B. & Chuong, C.-M. Early Events in Skin Appendage Formation: Induction of Epithelial Placodes and Condensation of Dermal Mesenchyme. J Invest Derm Symp P 4, 302–306 (1999).

31. Fuchs, E. Scratching the surface of skin development. Nature 445, 834–842 (2007).

32. Michon, F., Forest, L., Collomb, E., Demongeot, J. & Dhouailly, D. BMP2 and BMP7 play antagonistic roles in feather induction. Development 135, 2797–2805 (2008).

33. Palmquist, K. H. et al. Reciprocal cell-ECM dynamics generate supracellular fluidity underlying spontaneous follicle patterning. Cell 185, 1960–1973.e11 (2022).

34. Jensen, U. B., Lowell, S. & Watt, F. M. The spatial relationship between stem cells and their progeny in the basal layer of human epidermis: a new view based on whole-mount labelling and lineage analysis. Development 126, 2409–2418 (1999).

35. Kobayashi, Y. et al. Interplay between epidermal stem cell dynamics and dermal deformation. Npj Comput Mater 4, 45 (2018).

36. Viswanathan, P. et al. Mimicking the topography of the epidermal–dermal interface with elastomer substrates. Integr Biol 8, 21–29 (2015).

37. Mobasseri, S. A. et al. Patterning of human epidermal stem cells on undulating elastomer substrates reflects differences in cell stiffness. Acta Biomater 87, 256–264 (2019).

38. Marescal, O. & Cheeseman, I. M. Cellular Mechanisms and Regulation of Quiescence. Dev. Cell 55, 259–271 (2020).

39. Lång, E. et al. Coordinated collective migration and asymmetric cell division in confluent human keratinocytes without wounding. Nat Commun 9, 3665 (2018).

40. Lång, E., et al. Mechanical coupling of supracellular stress amplification and tissue fluidization during exit from quiescence. Proc. Natl. Acad. Sci. 119, e2201328119 (2022).

41. Széll, M. et al. Proliferating Keratinocytes Are Putative Sources of the Psoriasis Susceptibility-Related EDA+(Extra Domain A of Fibronectin) Oncofetal Fibronectin. J. Investig. Dermatol. 123, 537–546 (2004).

42. Mitra, M., Ho, L. D. & Coller, H. A. Cellular Quiescence, Methods and Protocols. Methods Mol. Biol. 1686, 27–47 (2017).

43. Coller, H. A., Sang, L. & Roberts, J. M. A New Description of Cellular Quiescence. PLoS Biol. 4, e83 (2006).

44. Li, B. et al. Autophagy mediates serum starvation-induced quiescence in nucleus pulposus stem cells by the regulation of P27. Stem Cell Res. Ther. 10, 118 (2019).

45. Gos, M. et al. Cellular quiescence induced by contact inhibition or serum withdrawal in C3H10T1/2 cells. Cell Prolif. 38, 107–116 (2005).

46. Barr, A. R. et al. DNA damage during S-phase mediates the proliferation-quiescence decision in the subsequent G1 via p21 expression. Nat. Commun. 8, 14728 (2017).

47. Henry, G., Li, W., Garner, W. & Woodley, D. T. Migration of human keratinocytes in plasma and serum and wound re-epithelialisation. Lancet 361, 574–576 (2003).

48. Bandyopadhyay, B. et al. A “traffic control” role for TGFβ3: orchestrating dermal and epidermal cell motility during wound healing. J. Cell Biol. 172, 1093–1105 (2006).

49. Rippon, M. G., Ousey, K. & Cutting, K. F. Wound healing and hyper-hydration: a counterintuitive model. J. Wound Care 25, 68–75 (2016).

50. Sweeney, I. R., Miraftab, M. & Collyer, G. A critical review of modern and emerging absorbent dressings used to treat exuding wounds. Int. Wound J. 9, 601–612 (2012).

51. Recho, P., Putelat, T. & Truskinovsky, L. Contraction-Driven Cell Motility. Phys. Rev. Lett. 111, 108102 (2013).

52. Keller, E. F. & Segel, L. A. Model for chemotaxis. J. Theor. Biol. 30, 225–234 (1971).

53. Dobin, A. et al. STAR: ultrafast universal RNA-seq aligner. Bioinformatics 29, 15–21 (2013).

54. Li, B. & Dewey, C. N. RSEM: accurate transcript quantification from RNA-Seq data with or without a reference genome. BMC Bioinformatics 12, 323–323 (2011).

55. Ge, S. X., Son, E. W. & Yao, R. iDEP: an integrated web application for differential expression and pathway analysis of RNA-Seq data. BMC Bioinformatics 19, 534 (2018).

