## Supplementary information for "Cell–cell adhesion drives patterning in stratified epithelia"

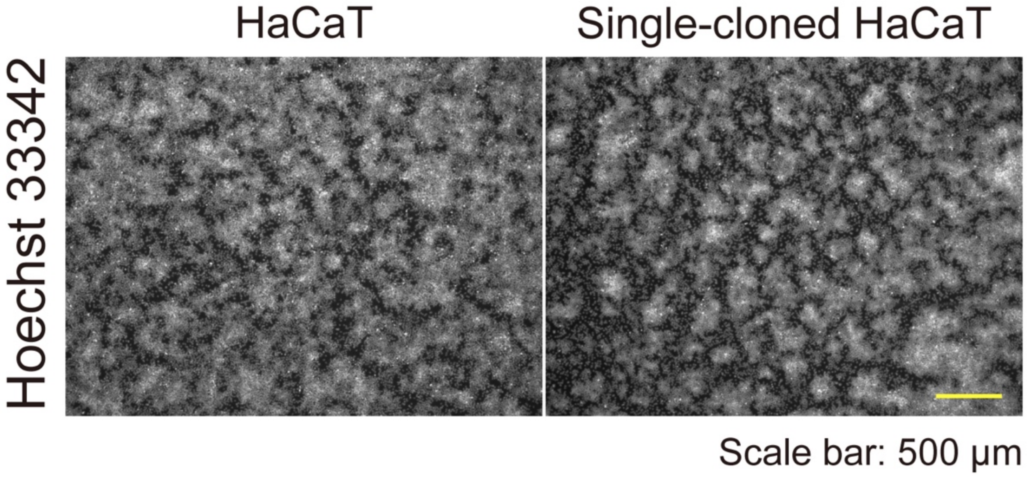


**Supplementary Figure 1. Parental and single-cell-cloned HaCaT cells.**

Immunofluorescent images of parental and single-cell-cloned HaCaT cells on day 4. Nuclei are labeled with Hoechst 33342. Scale bar: 500 µm.

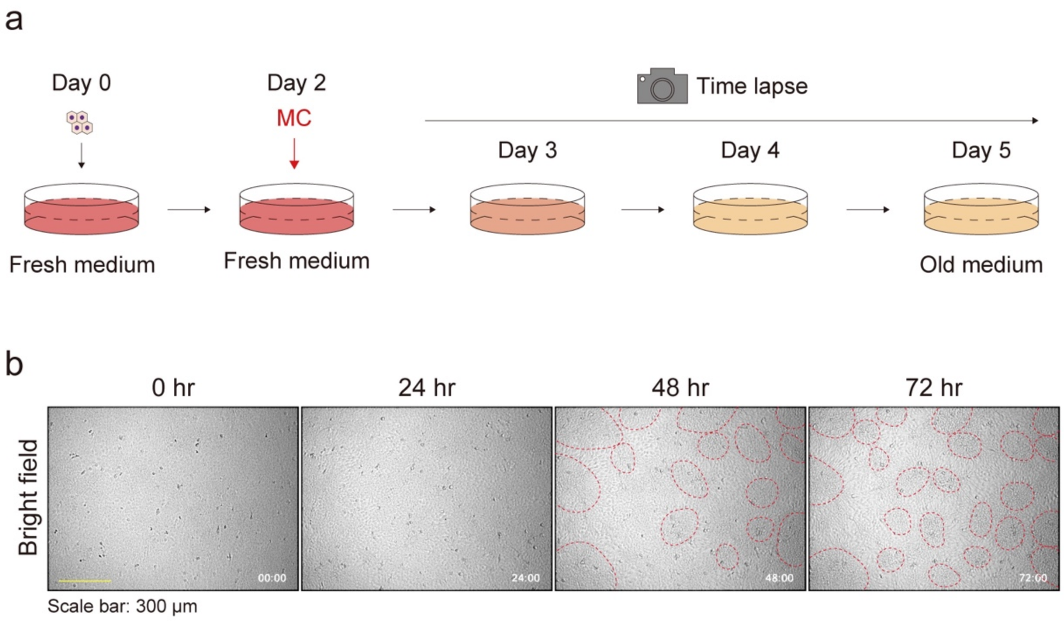


**Supplementary Figure 2. Keratinocyte patterns without medium change.**

**a** Schematic diagram of the time-lapse experiment for observing keratinocyte pattern formation. **b** Phase contrast time-lapse images of keratinocytes. Representative images are shown at 24-hour time intervals. Red dotted circles were manually added to highlight densely clustered areas. Each image is timestamped in hours and minutes. Scale bar: 300 µm.

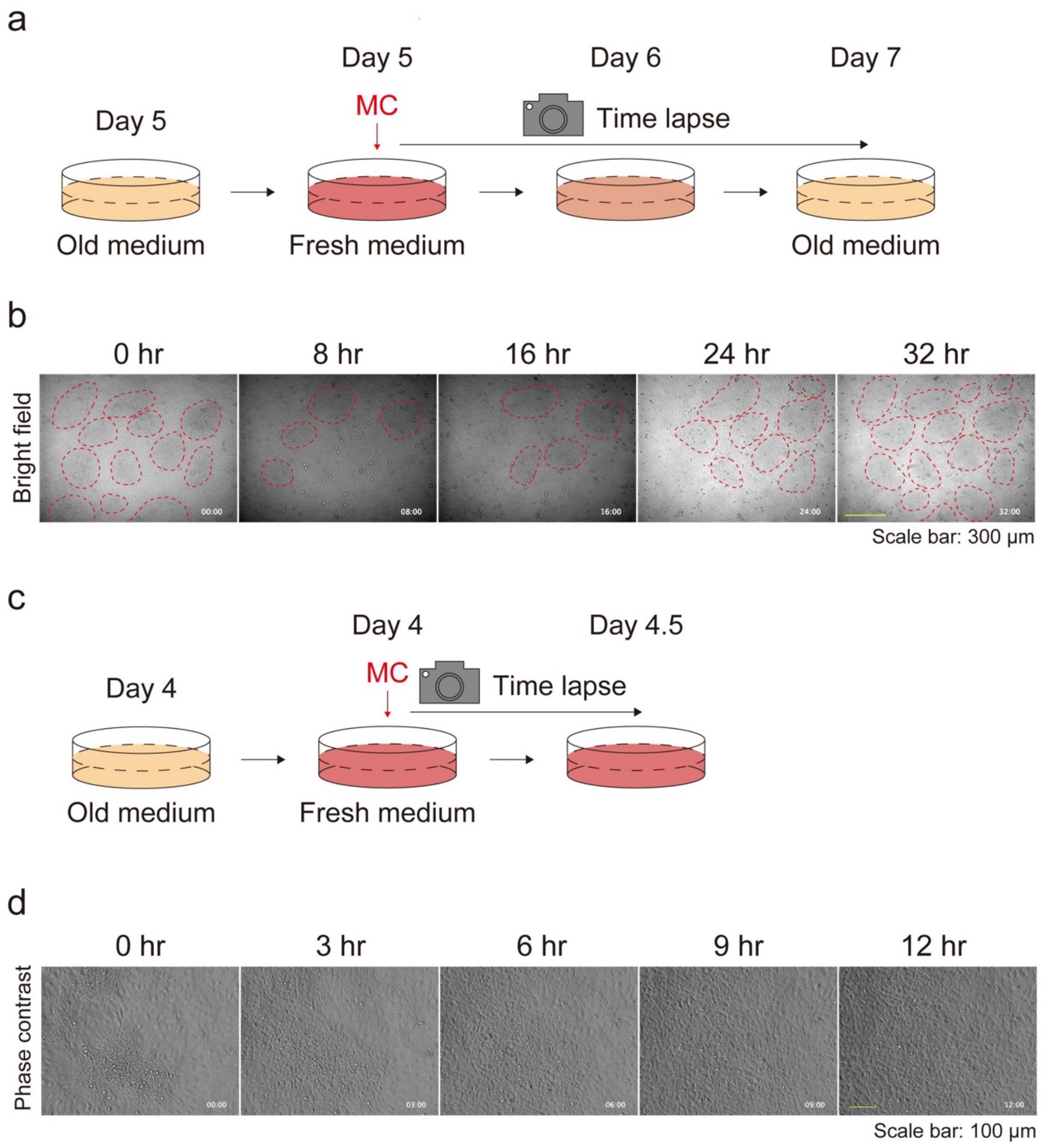


**Supplementary Figure 3. Keratinocyte patterns after medium replenishment.**

**a** Schematic diagram of time-lapse experiments for observing the keratinocyte patterns after medium replenishment. **b** Phase contrast time-lapse images of keratinocytes. Representative images are shown at 8-hour time intervals. Red dotted circles were manually added to identify densely clustered areas. Each image is timestamped in hours and minutes. Scale bar: 300 µm. **c** Schematic diagram of time-lapse experiments for high-magnification observation of the keratinocyte patterns after medium replenishment. **d** Phase contrast time-lapse high-magnification images of keratinocytes. Representative images are shown at 3-hour time intervals. Each image is timestamped in hours and minutes. Scale bar: 100 µm.

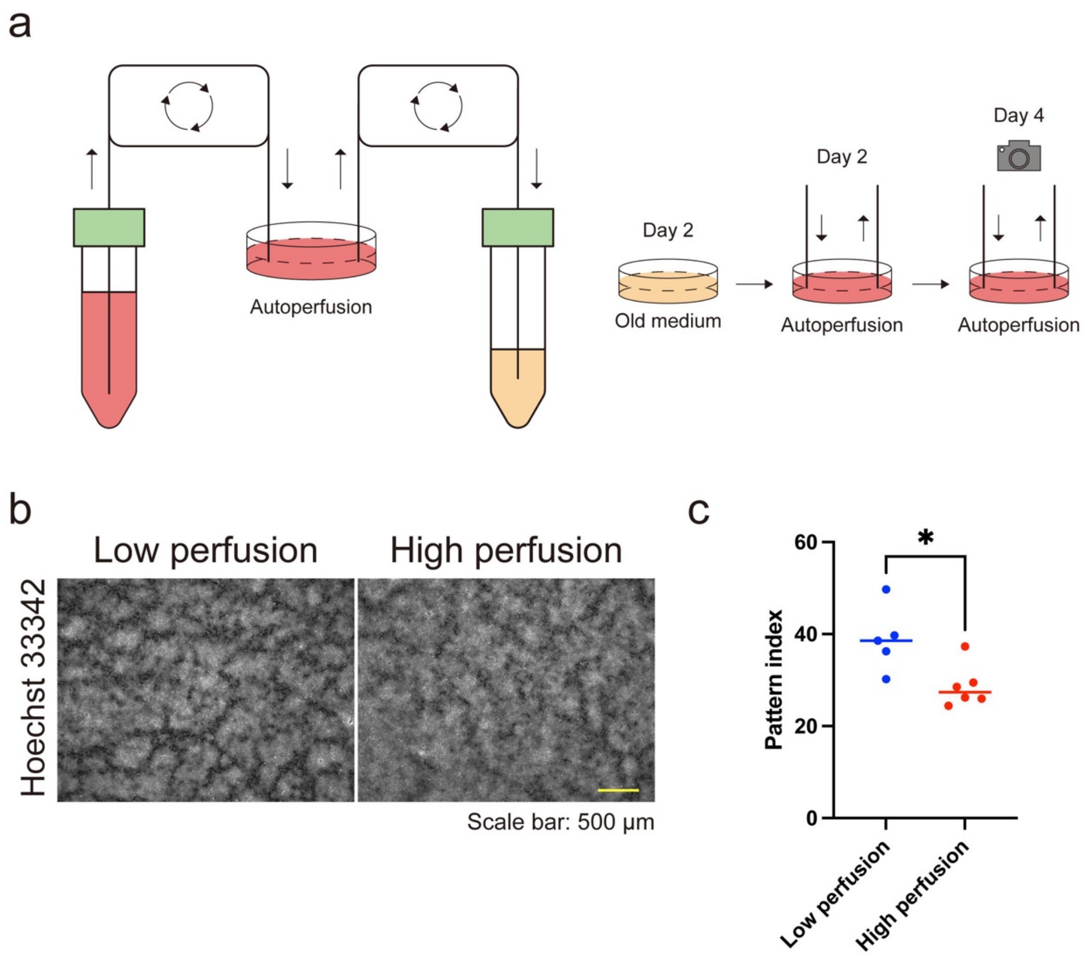


**Supplementary Figure 4. Keratinocyte patterns through autoperfusion.**

**a** Schematic diagram of the autoperfusion culture system and the autoperfusion culture experiment. **b** Immunofluorescent images of keratinocytes cultured under low and high perfusion rates on day 4. Nuclei are labeled with Hoechst 33342. Scale bar: 500 µm. **c** Pattern index for cultures with low and high perfusion rates. N = 5 for the culture group with the low perfusion rate and N = 6 for the culture group with the high perfusion rate. All data are presented as mean values and analyzed with two-tailed Mann–Whitney U tests. *, $P < 0.05$.

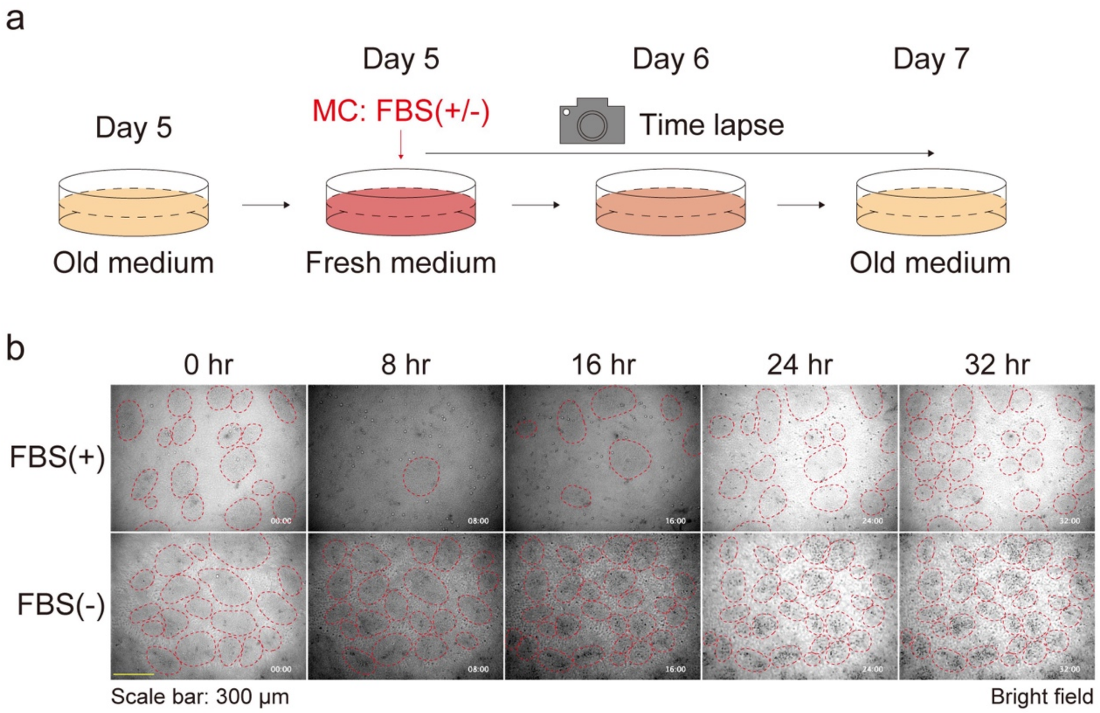


**Supplementary Figure 5. Keratinocyte patterns after medium replenishment with or without FBS.**

**a** Schematic diagram of time-lapse experiments for observing the keratinocyte patterns after medium replenishment with or without FBS. **b** Phase contrast time-lapse images of keratinocytes. Representative images are shown at 8-hour time intervals. Red dotted circles were manually added to identify high-density patterns. Each image is timestamped in hours and minutes. Scale bar: 300 μm.

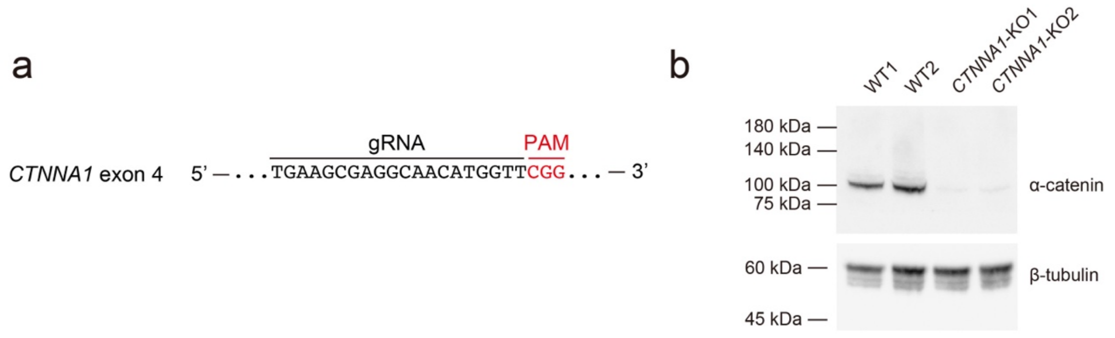


**Supplementary Figure 6. Establishment of *CTNNA1*-KO cells**

**a** The guide RNA design targeting exon 4 of the *CTNNA1* gene. **b** Western blot analysis using WT and *CTNNA1*-KO keratinocyte cell lysates. Immunoblotted by anti-α-catenin antibody and anti-β-tubulin antibody.

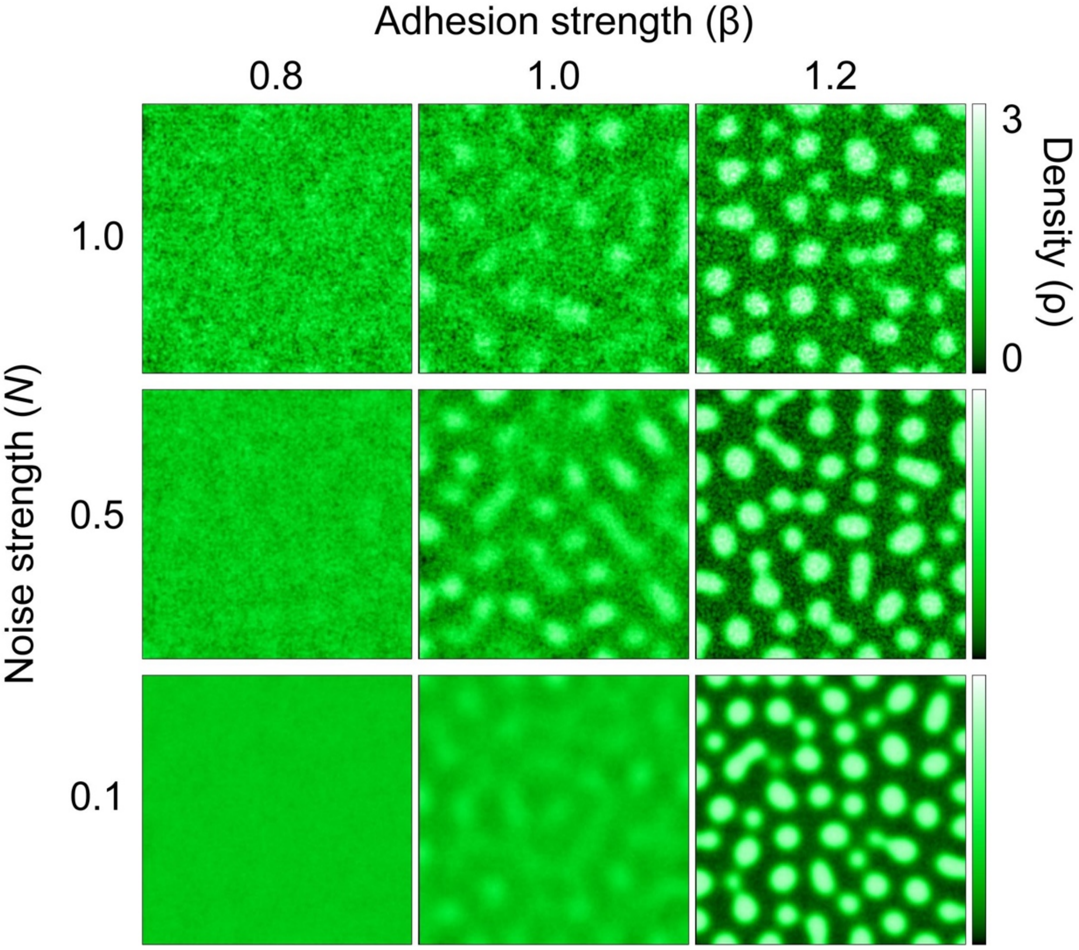


**Supplementary Figure 7. Mathematical modeling of a spatial density pattern using two variables: noise strength and adhesion strength.**

Simulated patterns of spatial cell distribution for varying adhesion strengths (β) and noise strengths (N), with time moment T [a.u.] = 120. Data for N = 0.5 are the same as in Figure 3.

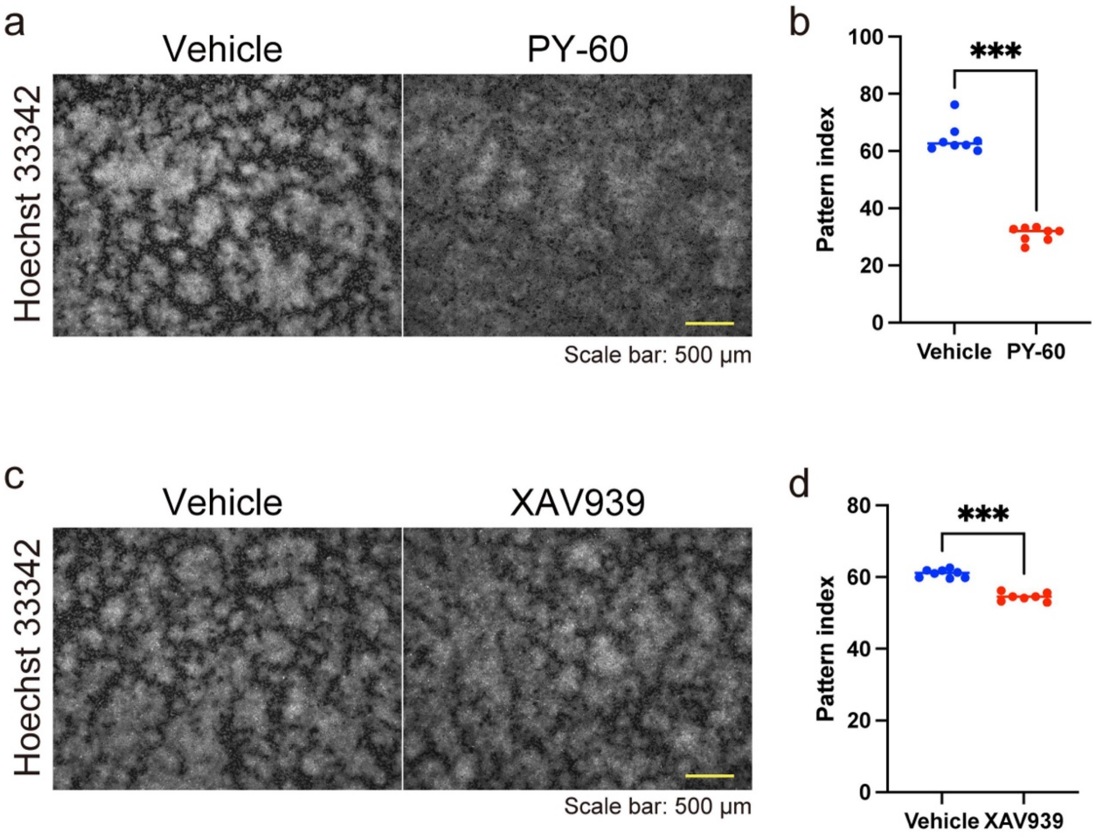


**Supplementary Figure 8. Patterns of keratinocytes treated with PY-60 or XAV939.**

**a** Immunofluorescent images of keratinocytes with or without PY-60 treatment. Nuclei are labeled with Hoechst 33342. Scale bar: 500 µm. **b** Pattern index with or without PY-80 treatment. N = 8 for each group. **c** Immunofluorescent images of keratinocytes with or without XAV939 treatment. Nuclei are labeled with Hoechst 33342. Scale bar: 500 µm. **d** Pattern index changes with or without XAV939 treatment. N = 8 for the vehicle group and N = 7 for the XAV939 treatment group. All data are presented as mean values and were analyzed with two-tailed Mann–Whitney U tests. ***, $P < 0.001$.

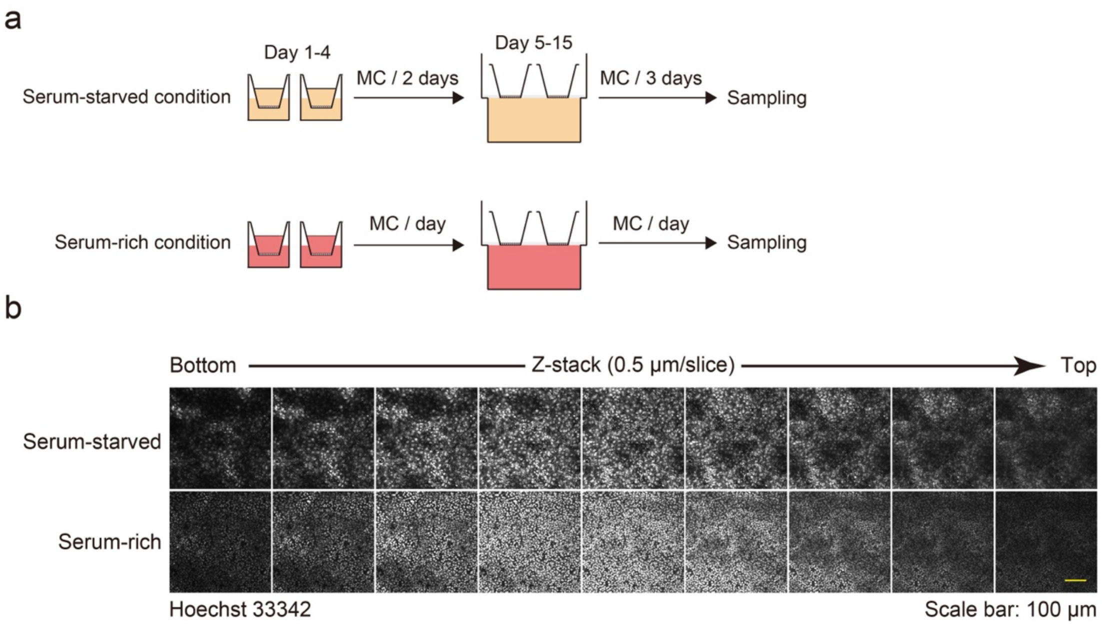


**Supplementary Figure 9. Keratinocyte patterns through 3D epidermis culture in serum-starved and serum-rich conditions.**

**a** Schematic diagram of 3D epidermis culture in serum-starved and serum-rich conditions. **b** Immunofluorescent images of keratinocytes through 3D culture. Nuclei are labeled with Hoechst 33342. Z-stack images are aligned from the bottom to the top. Z-stack depth: 0.5 μm/slice. Scale bar: 100 μm.
