## Supplementary material for "Cell–cell adhesion drives patterning in stratified epithelia": Description of additional supplementary files

**Supplementary Movie 1:**

A phase contrast time-lapse movie of keratinocytes without medium change (related to Supplementary Figure 2). Each image is timestamped in hours and minutes.

**Supplementary Movie 2:**

A phase contrast time-lapse movie of keratinocytes after medium replenishment (related to Supplementary Figure 3a, b). Each image is timestamped in hours and minutes.

**Supplementary Movie 3:**

A phase contrast time-lapse high-magnification movie of keratinocytes after medium replenishment (related to Supplementary Figure 3c, d). Each image is timestamped in hours and minutes.

**Supplementary Movie 4:**

A phase contrast time-lapse high-magnification movie of keratinocytes after medium replenishment with or without FBS (related to Supplementary Figure 5). Each image is timestamped in hours and minutes.

**Supplementary Movie 5:**

A simulation of the two-dimensional cell density model with the time moment T (related to Figure 3). The value of the adhesion strength (β) was 0.8. The color represents the local cell density ρ.

**Supplementary Movie 6:**

A simulation of the two-dimensional cell density model with the time moment T (related to Figure 3). The value of the adhesion strength (β) was 1.0. The color represents the local cell density ρ.

**Supplementary Movie 7:**

A simulation of the two-dimensional cell density model with the time moment T (related to Figure 3). The value of the adhesion strength (β) was 1.2. The color represents the local cell density ρ.
